# Meiotic drive is associated with sexual incompatibility in *Neurospora*

**DOI:** 10.1101/2020.11.27.400796

**Authors:** Aaron Vogan, Jesper Svedberg, Magdalena Grudzinska-Sterno, Hanna Johannesson

**Affiliations:** Department of Organismal Biology, Uppsala University, Sweden; Department of Biomolecular Engineering, Genomics Institute, UC Santa Cruz, USA

**Keywords:** Meiotic Drive, Bateson-Dobzhansky-Muller incompatibility, *Neurospora*, Speciation, Sexual incompatibility

## Abstract

Meiotic drive is the phenomenon whereby selfish elements bias their transmission to progeny at ratios above 50:50, violating Mendel’s law of equal segregation. The model fungus *Neurospora* carries three different meiotic drivers, called Spore killers. Two of these, *Sk-2* and *Sk-3*, are multilocus spore killers that constitute large haplotypes and are found in the species *N. intermedia*. Here we used molecular markers to determine that all *N. intermedia* isolates from New Zealand in fact belong to the sister species *N. metzenbergii*. Additionally, we use laboratory crosses to demonstrate that *Sk-2* and *Sk-3* are involved in sexual incompatibility between *N. intermedia* and *N. metzenbergii*.. Our experiments revealed that while crosses between these two species normally produced viable progeny at appreciable rates, when strains of *N. intermedia* carried *Sk-2* or *Sk-3* the proportion of viable progeny dropped substantially and in some crosses, no viable progeny were observed. Backcrossings supported that the incompatibility is tightly linked to the *Sk* haplotype. Finally, it appears that *Sk-2* and *Sk-3* have accumulated different incompatibility phenotypes when crossed with *N. metzenbergii* strains, consistent with their independent evolutionary history. This research illustrates how meiotic drive can contribute to reproductive isolation between populations, and thereby speciation.

## Introduction

Throughout the course of the last century, a significant amount of progress has been made in understanding how evolution creates diversity in the natural world and what factors drive speciation. It is understood that the phenomena controlling phenotypic divergence must coincide with changes at the genotypic level, and in terms of speciation, this divergence is often thought to be precipitated by so called Bateson-Dobzhansky-Muller (BDM) incompatibilities. In brief, the BDM model proposes that when two populations become isolated, changes at the genic level occur in such a way as for each population to evolve co-adapted alleles. When these two populations meet, the gene products from the different populations are no longer able to perform their function and attempts to mate result in inviable or sterile hybrid offspring. Despite the strong theoretical framework around the BDM model, empirical data on gene interactions that cause reproductive isolation between populations remain relatively rare (Maheshwari and Barbash 2011). As a result, the exact processes that lead to the formation of BMD incompatibilities are also poorly understood.

Meiotic drive is the phenomenon where a selfish genetic element is able to manipulate the sexual cycle of an organism such that it becomes over-represented in the offspring. Due to the mechanisms through which drivers cheat meiosis, there is often an associated penalty to fertility through the death of meiotic products, such as sperm or spores (Lindholm et al. 2016). Additionally, for many of the described cases of meiotic drivers, multiple genes are required for the drive to function, and these are kept in linkage through genomic inversions to suppress recombination (Presgraves 2009), which effectively isolates the driving haplotype from the non-driving version and can lead to the accumulation of linked deleterious mutations (Dyer et al. 2007). These regions of suppressed recombination are often large and encompass many genes not related to drive, which sets the stage for BDM incompatibilities to form. In agreement with this expectation, many cases have been described where individuals carrying meiotic drive elements show reproductive incompatibilities with non-driving individuals (Wilkinson and Fry 2001; Phadnis and Orr 2009; Zhang et al. 2015). Such incompatibilities may ultimately result in the inability of a driver to sweep to fixation in a population, as in order for driving alleles to reap the benefits of cheating meiosis, they must reproduce with naïve individuals. Alternatively, the reproductive isolation caused by the drive, could effectively split the population and result in speciation.

The filamentous fungus *Neurospora crassa*, and other closely related *Neurospora* species have been developed as a model system for comparisons of species recognition criteria and for the study of the evolution of reproductive isolation between species (Dettman et al. 2003a,b, 2006; Menkis et al. 2009; Villalta et al. 2009; Turner et al. 2011). They also harbour meiotic drivers. The three meiotic drivers known in *Neurospora* are *Sk-1, Sk-2*, and *Sk-3* and these function by means of spore killing, whereby spores that carry the driving element will kill their sibling spores that do not carry the driver. Spore killing has a very obvious phenotype in *Neurospora*. Following meiosis, one round of mitosis occurs to produce eight sexual spores (ascospores). These spores are packaged together in an ascus to form a single row. When spore killing occurs, four viable, black ascospores are observed together with four small aborted white ascospores in each ascus (Turner and Perkins 1979). *Sk-2* and *Sk-3* are multilocus drivers that were both discovered in the species *Neurospora intermedia*. In this species, the killing locus is linked to the resistance locus in a haplotype that extends across a 30 cM region surrounding the centromere on chromosome 3, resulting in a region of linkage covering roughly 400 genes (Svedberg et al. 2018). The two spore killers are mutual killers, meaning they produce no viable spores when crossed to each other, which is in contrast to crosses between two *Sk-2* strains or two *Sk-3* strains, which results in eight viable spores, i.e., no spore killing. Despite the fact that *Sk-2* and *Sk-3* utilise the same resistance gene and occupy a similar region of chromosome 3 (Hammond et al. 2012), they do not share the same killing gene and appear to have had a long history of evolutionary separation (Svedberg et al. 2018; Rhoades et al. 2019).

Thousands of wild isolates of *N. intermedia* have been collected from around the world and many of these have been evaluated by crosses to tester strains for the presence of *Sk-2* or *Sk-3*. Only four strains have been discovered that carry the *Sk-2* drive, and just one strain with *Sk-3*, but numerous strains have been isolated which are resistant to either *Sk-2, Sk-3*, or both. It has been suggested that the drivers may have had a large influence on the historic population structure of *N. intermedia*, but that widespread resistance has led to the recent decline of the drivers (Turner 2001). Genomic analyses suggest that the *Sk-2* haplotype diverged from other *N. intermedia* at least 250 thousand years ago, after the split of *N. intermedia* from its closest relative, *N. metzenbergii* (Svedberg et al. 2018). These results support a historically higher population size for *Sk-2*, as it is unlikely to remain at the present day frequencies for such a long period of time.

In an investigation of the geographic distribution of *Neurospora* spore killers, (Turner 2001) reported that strains of *N. intermedia* from Mexico and New Zealand (NZ) do not produce viable ascospores when mated to strains carrying *Sk-2* or *Sk-3* alleles. Additionally, introgressing the driver into a wild NZ genomic background failed to improve the number of viable ascospores in subsequent matings (Turner 2001), implying that reproductive isolation may be linked to the drivers themselves. Subsequent analyses of species diversity and classification in *Neurospora*, using mating behaviour and molecular markers resulted in the reclassification of many Mexican *N. intermedia* strains to a distinct species named *N. metzenbergii* (Dettman et al. 2003a,b; Villalta et al. 2009). *In this study, we hypothesise that the reproductive barrier observed by Turner (2001) relates to BDM incompatibilities between N. metzenbergii* specific alleles and loci linked to the *Sk* haplotypes. To test this hypothesis, we first extended the revision of *Neurospora* species classification initiated by Dettman et al. (2003a) and (Villalta et al. 2009) by determining the phylogenetic relationship of all the Mexican and NZ strains historically assigned to *N. intermedia* and show that, indeed, all of these strains belong to *N. metzenbergii*. Subsequently, by analyzing reproductive success in laboratory crosses, we verify that *N. metzenbergii* strains display a higher degree of reproductive isolation with *N. intermedia* strains carrying *Sk-2* or *Sk-3* than with those that do not. We found that the mechanism of reproductive isolation appears to differ between *Sk-2* and *Sk-3*, suggesting that the separate drivers have captured unique incompatibility factors during their independent evolutionary separation. We conclude that meiotic drive is specifically associated with strong reproductive isolation between strains of *Neurospora* and discuss the role that meiotic drive can play in speciation.

## Methods

### Strains

*Neurospora* strains used in this study were acquired from the Fungal Genetics Stock Centre (FGSC), Kansas State University. All strains annotated as *N. metzenbergii* (17) in the FGSC collection were included in the analysis, as well as all strains annotated as *N. intermedia* that were collected from New Zealand (11) or Mexico (1). Additional strains of *Neurospora* (54) were chosen to represent the diversity of the species based on Dettman et al. (2003a). We also included in our study strains of *N. intermedia* carrying either *Sk-2* or *Sk-3*. Note that for strain 7427 and 7428 (which contain the *Sk-2* allele) and strains 3193 and 3194 (containing the *Sk-3* allele), the original strains that carried the spore killer haplotype are not available from the FGSC. Instead, these *Sk-2* strains are F1 progeny of the wild spore killer strain and either of the *N. intermedia* tester strains 1766 or 1767, and the *Sk-3* strains are the result of three backcrosses from the original wild spore killer strains to the 1767 strain. Strain 7426 represents a wild strain that carries *Sk-2*, which was isolated at a later date. All strains used for mating assays are referred to by their FGSC ID number and are listed in **Table 1**. Note that 8761 and 8762 are single conidial isolates derived from 1766 and 1767, respectively. Locations of strains from the Perkin’s collection were taken from the FGSC (www.FGSC.net) and converted to global coordinates using the R package ggmap (Kahle and Wickham 2013).

**Table 1.** Strains used for crosses in this study

| Strain (FGSC) | Mating Type | Locale | Species |
| --- | --- | --- | --- |
| 5116 | A | New Zealand | *metzenbergii*<sup>a</sup> |
| 5117 | a | New Zealand | *metzenbergii*<sup>a</sup> |
| 5118 | a | New Zealand | *metzenbergii*<sup>a</sup> |
| 5119 | a | New Zealand | *metzenbergii*<sup>a</sup> |
| 1520 | A | New Zealand | *metzenbergii*<sup>a</sup> |
| 1521 | a | New Zealand | *metzenbergii*<sup>a</sup> |
| 1522 | A | New Zealand | *metzenbergii*<sup>a</sup> |
| 6795 | A | New Zealand | *metzenbergii*<sup>a</sup> |
| 6796 | a | New Zealand | *metzenbergii*<sup>a</sup> |
| 7829 | A | New Zealand | *metzenbergii*<sup>a</sup> |
| 7830 | a | New Zealand | *metzenbergii*<sup>a</sup> |
| 8846 | a | Mexico | *metzenbergii* |
| 8847 | A | Mexico | *metzenbergii* |
| 8849 | a | Mexico | *metzenbergii* |
| 8852 | a | Mexico | *metzenbergii* |
| 8853 | a | Mexico | *metzenbergii* |
| 8880 | A | Madagascar | *metzenbergii* |
| 8881 | a | Madagascar | *metzenbergii* |
| 10395 | a | Mexico | *metzenbergii* |
| 10396 | a | Mexico | *metzenbergii* |
| 10397 | a | Mexico | *metzenbergii* |
| 10399 | a | Haiti | *metzenbergii* |
| 10400 | a | Mexico | *metzenbergii* |
| 10401 | A | Mexico | *metzenbergii* |
| 10410 | a | Mexico | *metzenbergii* |
| 10411 | A | Mexico | *metzenbergii* |
| 10412 | a | Mexico | *metzenbergii* |
| 10413 | A | Mexico | *metzenbergii* |
| 8761 (1766) | A | Taiwan | *intermedia* |
| 8762 (1767) | a | Taiwan | *intermedia* |
| 7402 | a | Laboratory | *intermedia (Sk-2)* |
| 7427 | A | Laboratory | *intermedia (Sk-2)* |
| 7426 | A | Borneo | *intermedia (Sk-2)* |
| 7428 | a | Laboratory | *intermedia (Sk-2)* |
| 7429 | A | Laboratory | *intermedia (Sk-2)* |
| 3193 | A | Laboratory | *intermedia (Sk-3)* |
| 3194 | a | laboratory | *intermedia (Sk-3)* |
| 6263 | a | Ivory Coast | *intermedia* |
| 2316 | A | USA (Florida) | *intermedia* |
| 8795 | a | Papua New Guinea | *intermedia* |
| 2365 | a | USA (Hawaii) | *intermedia* |
| TSk2-Bc4 | A | Laboratory | *intermedia (Sk-2)* |
| TSk3-Bc4 | A | Laboratory | *intermedia (Sk-3)* |

<sup>a</sup>Previously classified as *N. intermedia*

### Molecular markers and phylogenetic analyses

We PCR-amplified and Sanger sequenced the markers TMI, TML, DMG, and QMA according to the protocols given in (Dettman et al. 2003a), to assign the investigated strains to species level groupings. These markers contain microsatellite sequences which were removed prior to alignment, and hence only the flanking sequences were aligned and analyzed, as in Dettman et al. (2003a). When comparing our sequencing results of a number of reference strains to those sequences generated by Dettman et al. (2003a) and subsequently used by Villalta et al (2009), we discovered a large number of cytosines in the Dettman et al. (2003a) sequences that are absent from ours, as well as from sequences from Corcoran et al. (2014) and Svedberg et al. (2018). Due to these discrepancies, we only used marker sequences that we generated ourselves in this study, or that were generated by Corcoran et al. (2014) (**Supplementary Table 1**). Sequences with the microsatellite portions removed were concatenated and aligned with the CLC sequence viewer v 7.7 (https://digitalinsights.qiagen.com/), followed by manual curation. We performed a Maximum Likelihood analysis using IQ-TREE v 2.0 (Minh et al. 2020) with the default parameters and 100 standard nonparametric bootstraps. Sequences are deposited in Genbank with accession numbers MW034381-MW034436.

Additionally, we performed a phylogenomic analysis based on SNP data from 40 strains distributed over eight species of *Neurospora* (**Supplementary Table 1**). Genomic data for these strains had been collected in previous studies (Corcoran et al. 2014; Svedberg et al. 2018) and SNPs were called against the *N. intermedia* strain 8807. For details on SNP calling, see (Svedberg et al. 2018). Whole chromosome phylogenies were inferred using RAxML (Stamatakis 2014) for six of the seven chromosomes. Chromosome 3 was excluded to remove the conflicting phylogenetic signal of the non-recombining spore killer region compared to the rest of the genome, even though that only affects the placement of strains within *N. intermedia*. The six phylogenetic trees were then merged into a consensus network with 20% pruning, using Splitstree (Huson and Bryant 2006).

### Crossings

All crosses were conducted in 10 × 100 mm glass culture tubes containing 1 mL of liquid synthetic crossing (SC) media (Westergaard and Mitchell 1947) with no added sucrose. A small strip of filter paper was added to the tubes before autoclaving. *N. intermedia* strains were used as females for all hybrid crosses: they were inoculated into the tubes and allowed to establish and produce protoperithecia (immature fruiting bodies) over 3 days at room temperature in complete darkness. Conidia (asexual spores that also function as fertilizing agents) from strains used as males in the crosses were then used to fertilize the protoperithecia and after fertilization, the cultures were incubated at 25°C in a 12h light/dark cycle. After 2 - 3 weeks mating had taken place for sexually compatible crosses, perithecia (mature fruiting bodies) were formed and ascospores had been shot, as evidenced by empty perithecia. To generate offspring from crosses, ascospores were harvested from the walls of the culture tubes using a sterile loop and spread onto the surface of water agar plates. These plates were incubated in a water bath at 60°C for 1 hour to induce germination as *Neurospora* ascospores require a heat treatment in order to germinate. This treatment kills any contaminating asexual spores (conidia) or mycelia, but will also kill non-pigmented ascospores. Germinated spores were picked from the water agar and placed into sterile glass culture tubes with Vogel’s media (Vogel 1956) to establish cultures.

### Estimation of reproductive success

To measure the reproductive success of crosses between strains of *Neurospora* we evaluated the production of perithecia and estimated the proportion of shot black spores to unpigmented white/hyaline spores, following Dettman et al. (2003b). In brief, crosses were allowed to mate until all ascospores had been shot. The sides of the culture tubes were examined under a dissecting microscope at 250X magnification and the proportion of black ascospores were estimated. The results of crosses were scored on a scale from 0 - 6 following Dettman et al. (2003b), where 0 refers to crosses that were entirely sterile (no fruiting bodies produced) and 6 refers to a fully compatible cross (fruiting bodies were produced and shot many black ascospores). The presence of an ostiole was not recorded as a measure of reproductive compatibility, hence categories 1 and 2 used by Dettman et al (2003b) are merged in this study. For selected combinations we set up the crossings in triplicate to determine final values. To verify the accuracy of the method and obtain quantitative values, proportions of black spores were also estimated through perithecial dissections for a limited number of crosses. For this method, we investigated the crosses at an earlier developmental stage, prior to the ejection of ascospores. Mature perithecia were identified based on the presence of long extended ostioles and selected for dissection. Perithecia were dissected on glass slides in a drop of sterile water to obtain rosettes of asci. Only rosettes which contained at least some fully matured black spores were evaluated for the proportion of viable spores. Additionally, perithecia were harvested over a 1 week period to verify that spore production was evaluated at the optimal time, i.e., after most asci had matured, but before most of the ascospores had been shot.

## Results

### Reassigning the New Zealand *N. intermedia* strains to *N. metzenbergii*

The relationship of the NZ *N. intermedia* strains (**Table 1**) to other *Neurospora* strains was assessed by creating a maximum likelihood tree from four microsatellite markers (TMI, DMG, TML, and QMA) as previously established by Dettman et al. (2003a) and Villalta et al. (2009) to recognize species in *Neurospora*. This tree (**Figure 1**) reveals that all available strains from NZ annotated in the FGSC as *N. intermedia* group with *N. metzenbergii* and fall within two divergent clades: one which places the NZ strains with the previously known clade of *N. metzenbergii* strains (isolated from Mexico, Haiti, and Madagascar) with robust support (Dettman et al. 2003a; Villalta et al. 2009), and another which constitutes only NZ strains.

**Figure 1.**
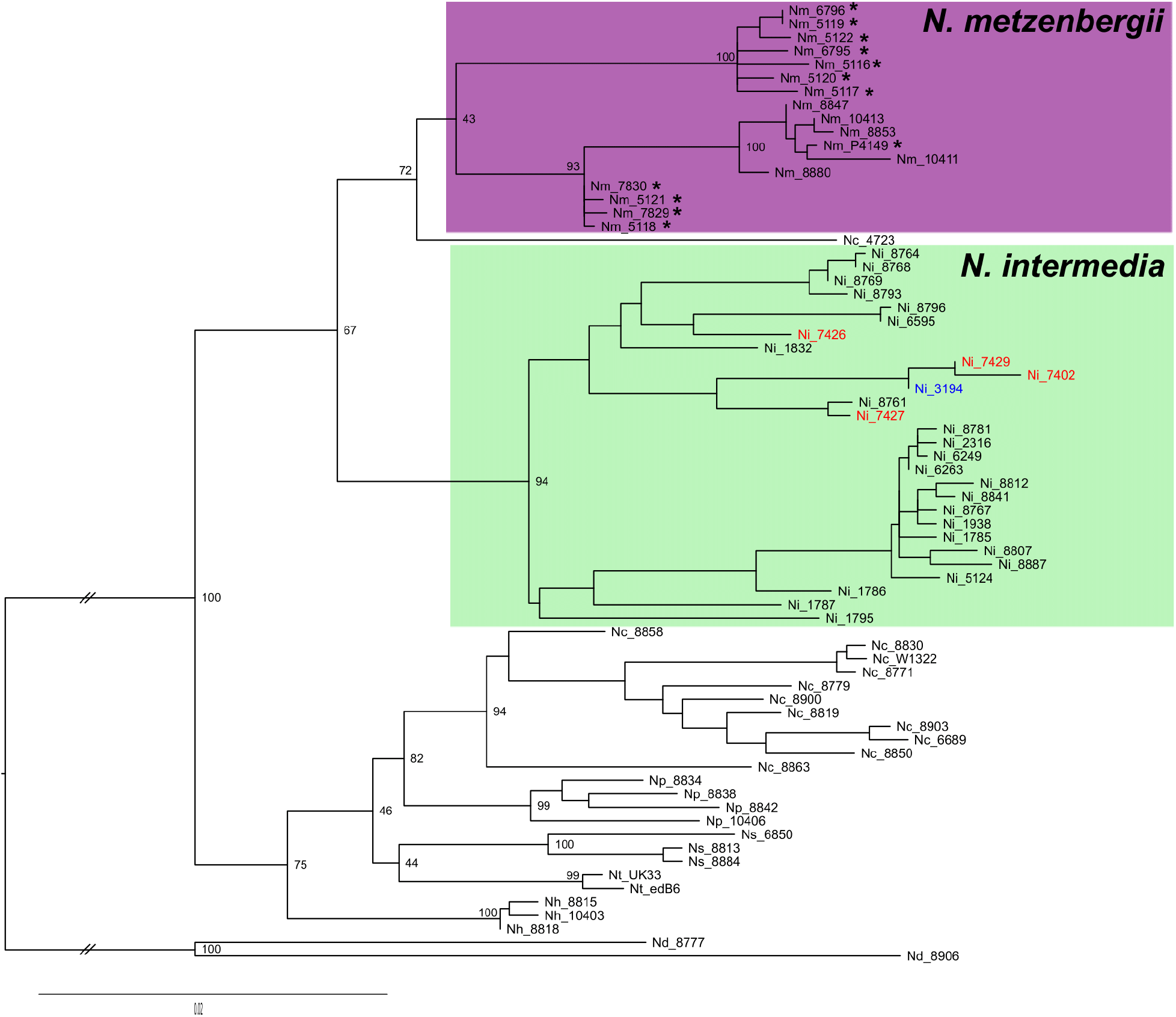
Maximum likelihood tree of 69 *Neurospora* strains, based on four genetic markers, TMI, TML, DMG, and QMA. *N. discreta* was set as the outgroup and the branch delineating the outgroup was shortened for illustration purposes. The spore killer strains are marked in red and blue for *Sk-2* and *Sk-3*, respectively. Sequences that were generated in this study are marked with an asterisk (*), all of which were formerly annotated as *N. intermedia*. The Nx prefixes prior to the FGSC numbers denote the species of the given strain.

However, note that the node supporting this sister relationship between the NZ strains and other *N. metzenbergii* strains has low support. In addition, we assessed one strain from Mexico annotated as *N. intermedia* in the FGSC (P4149), which also grouped well with the Mexican *N. metzenbergii* strains.

As the four marker tree did not fully resolve the relationships among the *Neurospora* species, we implemented a phylogenomic maximum likelihood analysis based on SNP data from whole genome assemblies from Illumina sequencing (**Supplementary Figure 1**). The overall groupings of strains within species agrees well with the marker tree and verifies that the NZ strains of *N. intermedia* belong to *N. metzenbergii*. Of note, the strain 4723, which has an unsupported position in the marker tree, falls outside all currently delimited species in the whole genome analysis. This strain was previously described as *N. crassa* based on sexual compatibility and was isolated along with another strain that was investigated in Dettman et al. (2003a) as a new candidate species, but ultimately rejected based on the marker tree. Our analysis from the genomic data suggests that it may indeed represent an undescribed species.

### Non-overlapping geographical distribution of *N. intermedia* and *N. metzenbergii*

In total, 3038 wild strains of *Neurospora* have been collected and were originally annotated as *N. intermedia* based on compatible mating to know reference strains, by David Perkins and others. By our extended revision of species recognition in *Neurospora*, as shown above, we reveal that all strains from New Zealand, Mexico and Madagascar, formerly referred to as *N. intermedia*, belong to *N. metzenbergii*. Based on this observation, it appears that while *N. intermedia* is plentiful in warmer climates (no isolates of *N. intermedia* were reported from Europe or northern North America), it never co-occurs with *N. metzenbergii* (**Figure 2**). This non-overlapping distribution is most striking around Mexico, where all isolates were determined as *N. metzenbergii*, while *N. intermedia* were found immediately to the north, in Texas, and to the south, in Honduras. The one exception is strain 10399 of *N. metzenbergii* which was isolated from Haiti alongside *N. intermedia* strains (**Figure 2**). It may be that these two species interact with competitive exclusion, but as strains were isolated at different times, temporal factors cannot be eliminated.

**Figure 2.**
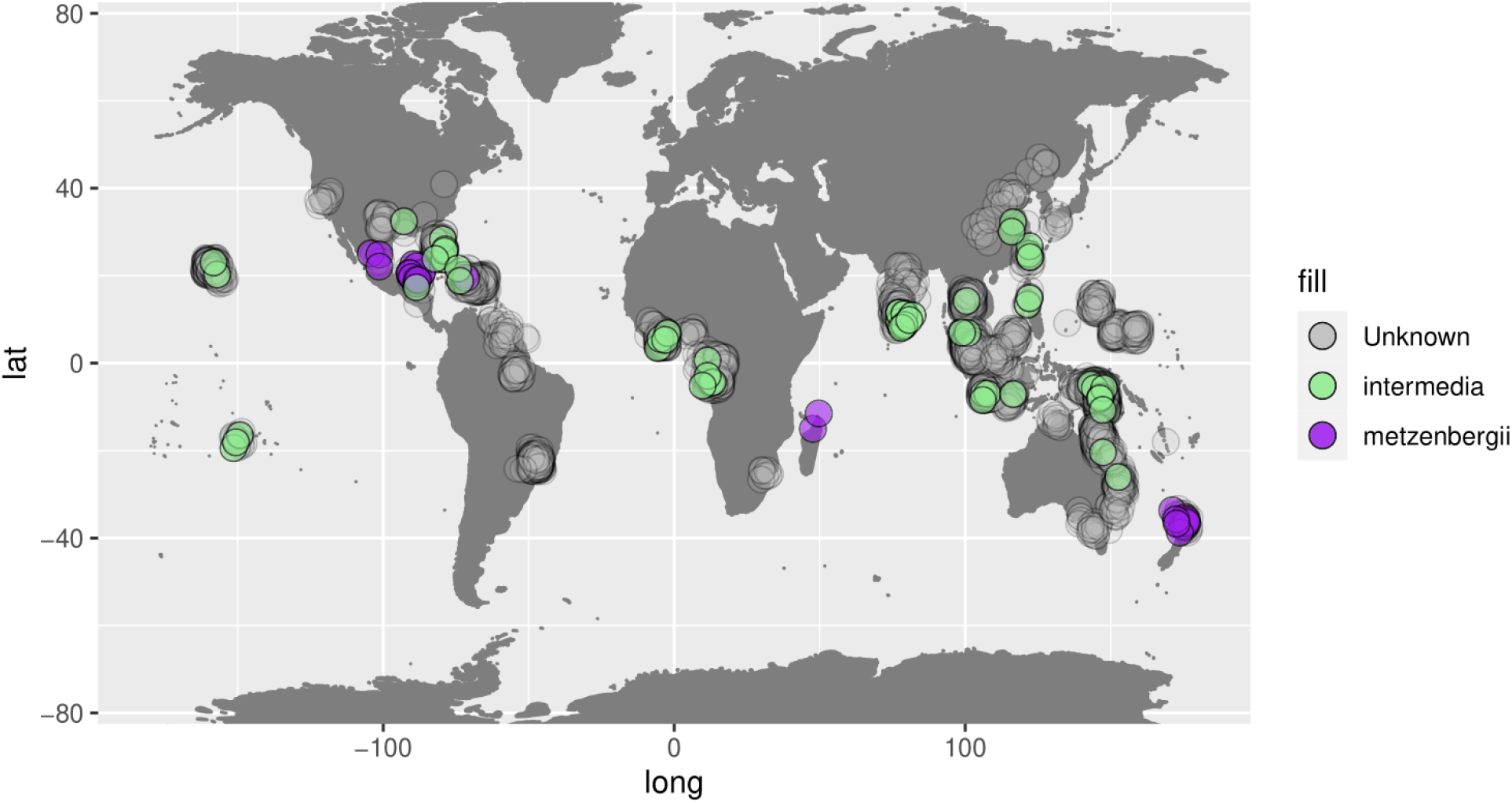
Global distribution of *N. intermedia* and *N. metzenbergii*. All strains from the Perkin’s collection which were determined to be *N. intermedia* through crossing to reference strains are plotted. The geographic origins of those which were confirmed as *N. intermedia* through molecular evidence are shown in green, those which were revealed to be *N. metzenbergii* are shown in purple. The “unknown” strains refer to isolates of *N. intermedia* with no molecular data.

### The spore killers of *N. intermedia* show higher incompatibility with *N. metzenbergii*

Previous work established that *N. metzenbergii* and *N. intermedia* are reproductively isolated, in that they produce a lower number of viable offspring when crossed to each other as opposed to crosses among strains of their own species (Dettman et al. 2003b; Villalta et al. 2009). We verified that the newly assigned NZ strains of *N. metzenbergii* show similar levels of reproductive isolation to *N. intermedia* by crossing representative strains of *N. metzenbergii* to standard tester strains of *N. intermedia* and among themselves. We see that, as expected, for both species the within-species crosses resulted in a high proportion of viable ascospores (>90%; **Supplementary Table 2**). Furthermore, most strains of *N. metzenbergii* were able to mate with the *N. intermedia* tester strains (1766 and 1767), but produced consistently low proportions of viable spores (60% - 80%). Notable exceptions to this range were the *N. metzenbergii* strain 10399 from Haiti, which produced even fewer viable spores (30% - 60%), and 7829 from New Zealand, which did not mate with any *N. intermedia* strains (**Supplementary Table 2**). Previous studies have also shown that some populations of *N. intermedia* have evolved strong pre-mating barriers with *N. crassa* (Turner et al. 2011). We investigated two strains from these populations here, 2316 and 6263, which revealed that these barriers may also prevent mating of these *N. intermedia* strains with strains of *N. metzenbergii* as successful matings were only achieved with one *N. metzenbergii* strain, 5120 (**Supplementary Table 2 and 3**).

In contrast to crosses with tester strains, crosses of the *N. metzenbergii* strains to the strains of *N. intermedia* that carry either *Sk-2* or *Sk-3* showed a dramatic reduction in the proportion of shot black spores. A drop of 50% in the proportion of viable spores is expected due to the spore killing itself, however many of the crosses resulted in nearly no black spores, or failure to produce perithecia at all, suggesting that there are strong incompatibilities linked to the spore killer region (**Supplementary Table 2**). To confirm that spore killing was occurring in these interspecific crosses, perithecia were dissected to screen for the standard 4:4 white to black spore pattern that should be observed. For all crosses between the spore killer strains and *N. metzenbergii* strains, the asci mostly contained either small aborted white spores, misshapen black spores, or white spores. In general, few asci were observed that could be properly evaluated for spore killing, but some did appear to have the standard 4:4 pattern, indicating that spore killing is active in the interspecific crosses.

Additionally, we used perithecial dissections of a subset of strains to quantify the degree of reproductive incompatibility (**Supplementary Table 3**). For all interspecific crosses, few asci were observed to have a full component of eight black ascospores. Nearly all asci contained at least one small aborted ascospore, and some asci appeared to be entirely aborted. Large proportions of partially pigmented ascospores or brown spores were also observed. Categorizing brown spores within asci is difficult, as these spores may have still developed into black spores prior to shooting. Additionally, some number of brown spores are viable and survive the heat treatment to germinate (Ho 1986). Due to these caveats, brown spores were considered viable in the data presented here. The perithecial dissections confirmed that some of the crosses of *N. metzenbergii* strains to *Sk-2* and *Sk-3* strains show a dramatic reduction in black spore production as compared to crosses with non-spore killer strains. Strains 10395 and 7830 did not exhibit a decrease in the proportion of black spores that was greater than the 50% expected from spore killing alone when crossed to the lab strains of *Sk-2* and *Sk-3*, however the methods used here may ignore completely aborted asci and so only quantify within ascus incompatibility. Strains 5119 and 8881 both show a considerable reduction in the proportion of black spores produced, with 8881 being the most severe. Crosses to the wild *Sk-2* strain 7426 were similar to the lab derived strains for 10395, but strains from NZ showed a large decrease in black spore production, suggesting that there may be additional incompatibilities in the non-spore killing region that have been lost in the lab derived strains (**Figure 3**).

**Figure 3.**
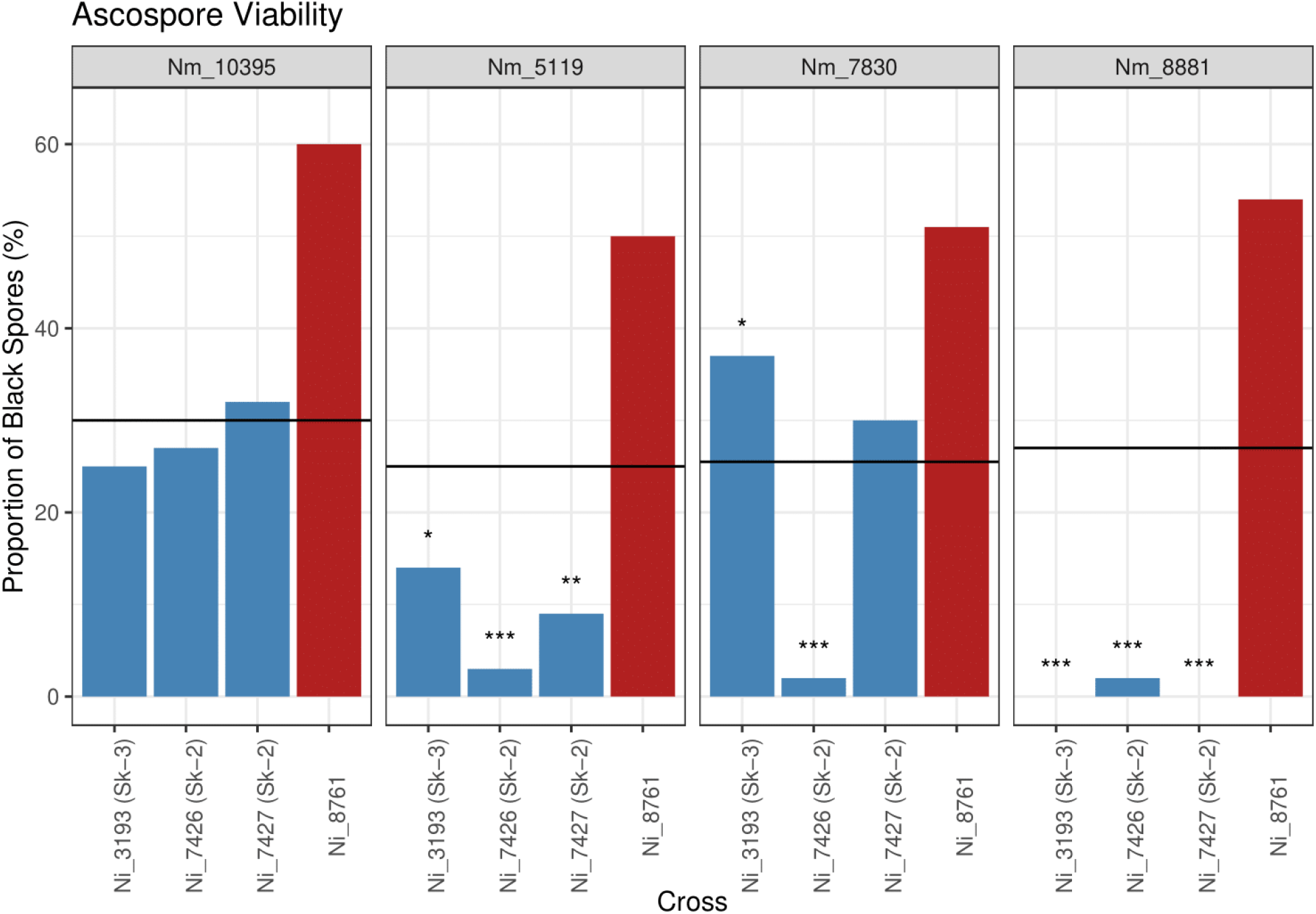
Proportion of black spores produced by crosses between four *N. metzenbergii* strains, 10395 (Mexico), 5119 (New Zealand), 7830 (New Zealand), and 8881 (Madagascar) and four *N. intermedia* strains 3193 (*Sk-3*), 7426 (*Sk-2*), 7427 (*Sk-2*), and 8761 (sensitive). Horizontal lines represent half the value of the cross to 8761 to the given *N. metzenbergii* strain as an expectation for a decrease in germination due to spore killing alone. Asterisks represent significant deviations from this expectation according to a chi square test (* < 0.5, ** < 0.01, *** < 0.001).

### Different incompatibility phenotypes of *Sk-2* and *Sk-3* with *N. metzenbergii*

A number of *N. metzenbergii* strains displayed different incompatibility phenotypes when crossed to *Sk-2* than with *Sk-3* despite the fact that the genomic background of the spore killer strains should be highly similar. With the Madagascar strains, 8880 and 8881, and strain 8846 (Mexico) no spores are shot when crossed to the Sk-2 strains, but many small aborted white spores are shot when crossed to the *Sk-3* strains, suggesting that there are different incompatibilities present within the different spore killer regions. To confirm that the spore killing region is specifically associated with the reproductive isolation observed between the Madagascar strains and the *Sk-2* and *Sk-3* strains, F1 progeny from crosses between 1767 and either 3193 (*Sk-3*) or 7427 (*Sk-2*) were backcrossed four times to 1766 to generate spore killer strains with a more isogenic background to 1766. Crosses to these 5x backcrossed strains showed identical phenotypes as to the parental spore killer strains, confirming the association of the phenomenon to the spore killer region. To verify these phenotypes, perithecial dissections were conducted. These confirmed that perithecia appear to be formed normally (ostioles are present) in crosses to both spore killers, but that in the *Sk-2* crosses, most asci abort at the spore-forming stage (**Figure 4A**), whereas in the *Sk-3* crosses, spores are formed, but nearly all are abortive (small and white) (**Figure 4B**). With crosses to *Sk-3*, occasionally viable spores can be found (**Figure 4B**), and these can germinate after heat treatment, showing they are fully viable. Given this low rate of viable spore production, it is likely due to rare recombination events within the spore killer region, which is hypothesised to occur occasionally (Svedberg et al. 2018)

**Figure 4.**
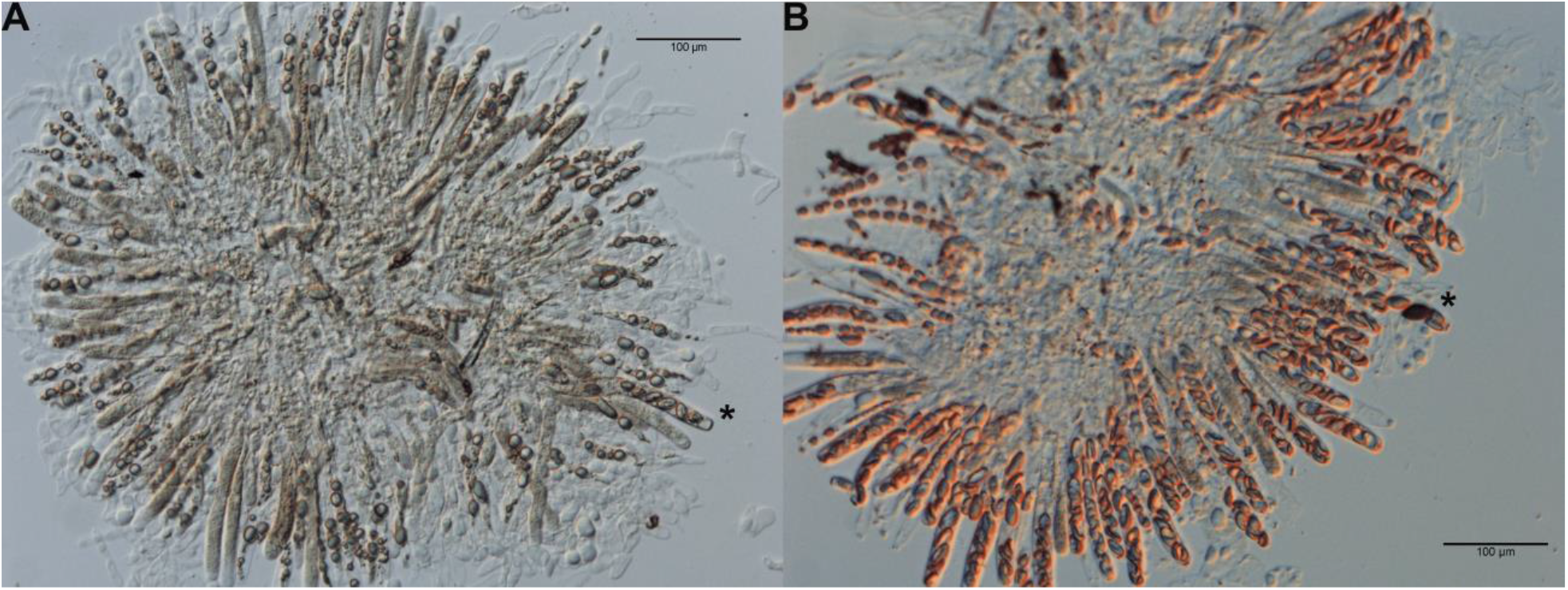
Rosettes of asci from crosses between *N. metzenbergii* 8881 and (**A**) the *Sk-2* backcross strains or (**B**) the *Sk-3* backcross strain. For crosses to *Sk-2*, nearly all asci are aborted without producing spores of any kind, but occasionally will produce an ascus with small aborted spores (*). For *Sk-3* crosses, asci contain only small aborted spores most of the time, but occasionally viable spores are found (*).

To investigate the incompatibilities further, hybrids were collected from crosses between the Madagascar strain 8880 and the *N. intermedia* strains 8761 and 8795, and crossed to the *Sk-2* and *Sk-3* strains. Most of the hybrids maintain the specific incompatibilities when crossed to *Sk-2* and *Sk-3* tester strains, however 8/36 recover some fertility. A few strains show differential recovery of fertility between *Sk-2* and *Sk-3*, in that some black spores were observed in crosses to the *Sk-2* strain, but not the *Sk-3* strain, or vice versa, supporting the hypothesis that the different phenotypes observed between *Sk-2* and *Sk-3* crosses represent different underlying incompatibilities.

## Discussion

The role that meiotic drivers, in particular sex-biased drives, can play in speciation processes has been under considerable debate, with opinions regarding the ability and importance of drivers to cause speciation varying wildly (Frank 1991; Coyne and Orr 1993; Zhang et al. 2015; Sweigart et al. 2019). Here we provide evidence that meiotic drive in *Neurospora* contributes to reproductive isolation, both directly and through the accumulation of linked incompatibilities, a process that is often overlooked in discussions of drive and speciation (Sweigart et al. 2019).

While the low frequency of both drive and full incompatibility suggests that the precise incompatibilities at play in *Sk-2* and *Sk-3* are unlikely to have been major contributors in the divergence of *N. metzenbergii* and *N. intermedia*, it nevertheless provides an example of a scenario where meiotic drive could directly drive speciation in other systems.

In order to understand what conditions may have led to the evolution of reproductive incompatibilities between *N. intermedia* and *N. metzenbergii*, it is important to determine what evolutionary forces were likely most dominant during their divergence. The delimitation between *N. intermedia* and *N. metzenbergii* aligns well with both the biological species concept and the phylogenetic species concept, in that molecular markers demonstrate their monophyly and mating assays show them to reproduce better with conspecifics than interspecifics (Dettman et al. 2003a,b). Here we have demonstrated that strains of *Neurospora* that were isolated from New Zealand, and previously designated as *N. intermedia*, in fact belong to *N. metzenbergii*, nearly doubling the number of known *N. metzenbergii* samples in collections and highlighting the unusual geographical distribution of the two species, which is consistent with them competing to the end of competitive exclusion. Together, these facts all point to the possibility that the *N. intermedia* and *N. metzenbergii* originally diverged in allopatrically (species with overlapping ranges tend to compete less (Letcher et al. 1994)), thereby fulfilling the main requirement for the formation of BDM incompatibilities.

We have provided evidence that additional reproductive barriers exist within the spore killer region of Sk-2 and Sk-3 strains of *N. intermedia* that inhibit the ability of said strains to hybridize with *N. metzenbergii*. This is partially driven by the 50% loss of spores due to spore killing, but the strains carrying *Sk-2* and *Sk-3* also exhibit a strong reduction in fertility with some strains of *N. metzenbergii* that go beyond what can be explained by the activity of the spore killer elements alone. It could be expected that the reason crosses between the spore killers and *N. metzenbergii* strains show a decrease in viable progeny is the production of unbalanced gametes as a result of the mismatched inversions, but as a reduced fertility is not observed between crosses of spore killer and sensitive strains within *N. intermedia*, it is unlikely to be a significant issue. Additionally, the fact that the strongest incompatibilities with *N. metzenbergii* strains appears to be polymorphic, further argues against a significant role of the inversions.The occurrence of polymorphic incompatibility factors between populations has been discussed as variable reproductive isolation (VRI), and has been observed in a variety of systems (Cutter 2012). Under a VRI scenario, these polymorphic alleles could originate either from ancestral, incomplete lineage sorting or as derived alleles within *N. metzenbergii*, but derived alleles are more likely to contribute to reproductive isolation (Orr 1995). Moreover, as only the killer locus allele is incompatible with these *N. metzenbergii* strains, the incompatible allele of *N. intermedia* must be located within the *Sk* haplotypes. Additionally, the differential phenotypes exhibited between *Sk-2* and *Sk-3* crosses suggest that discrete incompatibile loci of large effect exist within the *Sk-2* and *Sk-3* regions. Conversely, as *Sk-2* and *Sk-3* do not appear to share the same killer gene (Rhoades et al. 2019) it could be possible that the spore killing genes themselves are responsible for the incompatibility, as has been observed with one meiotic driver from *Drosophila* (Phadnis and Orr 2009).

The fact that incompatibilities are observed to accumulate in the non-recombining killer locus potentially provides deeper insights into the evolution of BDM incompatibilities. While the per base pair rate of divergent substitutions between *N. intermedia* and *N. metzenbergii* is higher in the *Sk* region than the rest of the genome, these still represent the minority of total divergent sites across the genome (10,345/191,911 substitutions) and the spore killer strains, in fact, have slightly fewer nonsynonymous substitutions in total than sensitive strains (59,077 vs. 63,279) (Svedberg et al. 2018). It is known that selection plays an important role in the accumulation of BDM incompatibilities and while a lot of attention has been payed to directional selection and stabilizing selection (Welch 2004; Fierst and Hansen 2010; Nosil and Schluter 2011), less has been given to decreased purifying selection. Additionally, a considerable amount of theory and modelling have discussed the role of deleterious recessive mutations linked to drivers or regions of reduced recombination in speciation (Navarro and Barton 2003; Welch 2004; Kirkpatrick and Barton 2006). We can be confident that deleterious recessive mutations are not a factor in *Sk-2* or *Sk-3* as crosses among killer strains of the same type show no decrease in the production of viable offspring. Nonetheless, the spore killer regions may be accumulating non-functional mutations due to the relaxed selection in the haplotype. Importantly, only non-functional mutations that can be compensated for by the genomic background of *N. intermedia* will persist, as others will be lethal. However, in the hybrid crosses to *N. metzenbergii*, alleles allowing for such compensation may, in some cases, not be present. So while phylogenomic analyses suggest that most sites in the *Sk* haplotype are diverging under neutral expectation (Svedberg et al. 2018) This scenario is similar to the “faster-X” theory of why sex chromosomes accumulate incompatibilities faster than autosomes (Mank et al. 2010; Lima 2014). These ideas should hold true for any genomic region experiencing suppressed recombination, such as inversions, and for this reason, have broader implications beyond meiotic drive.

Hybrid incompatibilities have been linked to meiotic drivers in numerous systems (Braidotti and Barlow 1997; Tao et al. 2001; Wilkinson et al. 2003; McDermott and Noor 2010), but the discussion of how they could contribute to speciation has largely centered around the coevolution of complex suppressor systems, and the divergence of those systems in disparate populations (Courret et al. 2019). This discussion may be largely irrelevant here as no suppressors are known to be involved in *Sk-2* or *Sk-3*, nor is there any evidence of other meiotic drivers within *N. metzenbergii*. With spore killing (and mechanistically similar meiotic drivers), if mutual killers, such as *Sk-2* and *Sk-3*, fix in different populations, then crosses between individuals of those populations will be entirely incompatible. Such a scenario appears to have arisen in the fission yeast, *Schizosaccharomyces pombe*, where the diverse group of *wtf* genes show unique distributions in individual lineages (Bravo Núñez et al. 2020). Mutual killing may be a common phenomenon as it is observed in three well described systems, *S. pombe, N. intermedia*, and for the *Spok* genes of *Podospora anserina* (Vogan et al. 2019). However, such a mechanism will only remain in place as long as the drive is active, if the driver reaches fixation in a given population, it may be prone to decay (as observed in both *S. pombe* and *P. anserina*) and the reproductive barrier that it caused would disappear. Therefore, cases where the driving mechanism and/or co-evolved suppressor genes implicitly cause reproductive isolation may not be expected to persist through time to impact speciation. However, if BDM incompatibilities have time to form during this phase, or as discussed above, are even more prone to formation in these regions, then drive could still contribute significantly to the reproductive isolation between populations, even if said drive then subsequently vanishes from the populations.

It should be noted that both the *Spok* and *wtf* genes are single gene drivers, which should be much less prone to the accumulation of linked incompatibilities illustrated here. Additionally, these small loci could be prone to cross species boundaries, as is similarly observed in *Drosophila* (Meiklejohn et al. 2018). In fact the third known spore killer in *Neurospora, Sk-1* from *N. sitophila*, is also a single gene driver and has also been observed to jump species boundaries (Svedberg J, Vogan AA, Rhoades NA, Sarmarajeewa D, Jacobson DJ, Lascoux M, Hammond TM, Johannesson H. 2020). In a scenario where reproductive isolation is caused by mutual killers, if an individual were able to acquire both drivers, it would be resistant to both drives and therefore viable with both species/populations. The selective advantage for such an individual could be quite strong, arguing against single gene drives as speciation genes. This implies that meiotic drivers that constitute large haplotypes may have greater impacts on population structure than single gene drives, and hence be observed more often during interspecies/population crossings. Therefore, although until recently, meiotic drives were thought to require multiple linked loci, it may be the case that single gene drivers are much more common, but are observed less due to minimal linked phenotypes.

## Acknowledgements

This work was supported by the European Research Council (ERC) grant ERC-2014-CoG (project 648143, SpoKiGen) and The Swedish Research Council to H.J. We thank the support given by the National Genomics Infrastructure (NGI) / Uppsala Genome center on massive parallel DNA sequencing. The computations were performed on resources provided by SNIC through Uppsala Multidisciplinary Center for Advanced Computational Science (UPPMAX) under the project SNIC 2017/1-567. We would also like to thank Dave Jacobson for the invaluable advice and data on the Perkin’s collection.

## Supplementary Figures

**Supplementary Figure 1**. Network based on maximum likelihood trees inferred from whole chromosome SNP data from six of the seven chromosomes.

## Supplementary Tables

**Supplementary Table 1**. Strains and molecular markers included in this study.

**Supplementary Table 2**. Proportion of shot black ascospores from laboratory crosses.

**Supplementary Table 3**. Proportion of black ascospores of laboratory crosses from dissected perithecia.

## References

Braidotti, G., and D. P. Barlow. 1997. Identification of a male meiosis-specific gene, Tcte2, which is differentially spliced in species that form sterile hybrids with laboratory mice and deleted in t chromosomes showing meiotic drive. Dev. Biol. 186:85–99.

Bravo Núñez, M. A., I. M. Sabbarini, M. T. Eickbush, Y. Liang, J. J. Lange, A. M. Kent, and S. E. Zanders. 2020. Dramatically diverse Schizosaccharomyces pombe wtf meiotic drivers all display high gamete-killing efficiency. PLoS Genet. 16:e1008350.

Corcoran, P., J. R. Dettman, Y. Sun, E. M. Luque, L. M. Corrochano, J. W. Taylor, M. Lascoux, and H. Johannesson. 2014. A global multilocus analysis of the model fungus Neurospora reveals a single recent origin of a novel genetic system. Mol. Phylogenet. Evol. 78:136–147.

Courret, C., C.-H. Chang, K. H.-C. Wei, C. Montchamp-Moreau, and A. M. Larracuente. 2019. Meiotic drive mechanisms: lessons from. Proc. Biol. Sci. 286:20191430.

Coyne, J. A., and H. A. Orr. 1993. Further evidence against meiotic-drive models of hybrid sterility. Evolution 47:685–687.

Cutter, A. D. 2012. The polymorphic prelude to Bateson–Dobzhansky–Muller incompatibilities.

Dettman, J. R., D. J. Jacobson, and J. W. Taylor. 2003a. A multilocus genealogical approach to phylogenetic species recognition in the model eukaryote Neurospora. Evolution 57:2703–2720.

Dettman, J. R., D. J. Jacobson, and J. W. Taylor. 2006. Multilocus sequence data reveal extensive phylogenetic species diversity within the Neurospora discreta complex. Mycologia 98:436–446.

Dettman, J. R., D. J. Jacobson, E. Turner, A. Pringle, and J. W. Taylor. 2003b. reproductive isolation and phylogenetic divergence in Neurospora: comparing methods of species recognition in a model eukaryote. Evolution 57:2721–2741.

Dyer, K. A., B. Charlesworth, and J. Jaenike. 2007. Chromosome-wide linkage disequilibrium as a consequence of meiotic drive. Proc. Natl. Acad. Sci. U. S. A. 104:1587–1592.

Fierst, J. L., and T. F. Hansen. 2010. Genetic architecture and postzygotic reproductive isolation: evolution of Bateson-Dobzhansky-Muller incompatibilities in a polygenic model. Evolution 64:675–693.

Frank, S. A. 1991. Divergence of Meiotic Drive-Suppression Systems as an Explanation for Sex-Biased Hybrid Sterility and Inviability.

Hammond, T. M., D. G. Rehard, H. Xiao, and P. K. T. Shiu. 2012. Molecular dissection of Neurospora Spore killer meiotic drive elements. Proc. Natl. Acad. Sci. U. S. A. 109:12093–12098.

Ho, C. C. 1986. Identity and characteristics of Neurospora intermedia responsible for oncom fermentation in Indonesia. Food Microbiol. 3:115–132.

Huson, D. H., and D. Bryant. 2006. Application of phylogenetic networks in evolutionary studies. Mol. Biol. Evol. 23:254–267.

Kahle, D., and H. Wickham. 2013. ggmap: Spatial Visualization with ggplot2. The R Journal. 5:144–161.

Kirkpatrick, M., and N. Barton. 2006. Chromosome inversions, local adaptation and speciation. Genetics 173:419–434.

Letcher, A. J., A. Purvis, S. Nee, and P. H. Harvey. 1994. Patterns of Overlap in the Geographic Ranges of Palearctic and British Mammals.

Lima, T. G. 2014. Higher levels of sex chromosome heteromorphism are associated with markedly stronger reproductive isolation. Nat. Commun. 5:4743.

Maheshwari, S., and D. A. Barbash. 2011. The Genetics of Hybrid Incompatibilities. Annu. Rev. Genet. 45:331–355.

Mank, J. E., B. Vicoso, S. Berlin, and B. Charlesworth. 2010. Effective population size and the Faster-X effect: empirical results and their interpretation. Evolution 64:663–674.

McDermott, S. R., and M. A. F. Noor. 2010. The role of meiotic drive in hybrid male sterility. Philos. Trans. R. Soc. Lond. B Biol. Sci. 365:1265–1272.

Meiklejohn, C. D., E. L. Landeen, K. E. Gordon, T. Rzatkiewicz, S. B. Kingan, A. J. Geneva, J. P. Vedanayagam, C. A. Muirhead, D. Garrigan, D. L. Stern, and D. C. Presgraves. 2018. Gene flow mediates the role of sex chromosome meiotic drive during complex speciation. Elife 7.

Menkis, A., E. Bastiaans, D. J. Jacobson, and H. Johannesson. 2009. Phylogenetic and biological species diversity within the Neurospora tetrasperma complex. J. Evol. Biol. 22:1923–1936.

Minh, B. Q., H. A. Schmidt, O. Chernomor, D. Schrempf, M. D. Woodhams, A. von Haeseler, and R. Lanfear. 2020. IQ-TREE 2: New Models and Efficient Methods for Phylogenetic Inference in the Genomic Era. Mol. Biol. Evol. 37:1530–1534.

Navarro, A., and N. H. Barton. 2003. Accumulating postzygotic isolation genes in parapatry: a new twist on chromosomal speciation. Evolution 57:447–459.

Nosil, P., and D. Schluter. 2011. The genes underlying the process of speciation.

Orr, H. A. 1995. The population genetics of speciation: the evolution of hybrid incompatibilities. Genetics 139:1805–1813.

Phadnis, N., and H. A. Orr. 2009. A single gene causes both male sterility and segregation distortion in Drosophila hybrids. Science 323:376–379.

Presgraves, D. 2009. Drive and sperm. Pp. 471–506 in Sperm Biology.

Rhoades, N. A., A. M. Harvey, D. A. Samarajeewa, J. Svedberg, A. Yusifov, A. Abusharekh, P. Manitchotpisit, D. W. Brown, K. J. Sharp, D. G. Rehard, J. Peters, X. Ostolaza-Maldonado, J. Stephenson, P. K. T. Shiu, H. Johannesson, and T. M. Hammond. 2019. Identification of rfk-1, a Meiotic Driver Undergoing RNA Editing in Neurospora. Genetics 212: 93-110.

Stamatakis, A. 2014. RAxML version 8: a tool for phylogenetic analysis and post-analysis of large phylogenies. Bioinformatics 30:1312–1313.

Svedberg, J., S. Hosseini, J. Chen, A. A. Vogan, I. Mozgova, L. Hennig, P. Manitchotpisit, A. Abusharekh, T. M. Hammond, M. Lascoux, and H. Johannesson. 2018. Convergent evolution of complex genomic rearrangements in two fungal meiotic drive elements. Nat. Commun. 9:4242.

Svedberg J, Vogan AA, Rhoades NA, Sarmarajeewa D, Jacobson DJ, Lascoux M, Hammond TM, Johannesson H. 2020. An introgressed gene causes meiotic drive in Neurospora sitophila. bioRxiv.

Sweigart, A. L., Y. Brandvain, and L. Fishman. 2019. Making a Murderer: The Evolutionary Framing of Hybrid Gamete-Killers. Trends Genet. 35:245–252.

Tao, Y., D. L. Hartl, and C. C. Laurie. 2001. Sex-ratio segregation distortion associated with reproductive isolation in Drosophila. Proc. Natl. Acad. Sci. U. S. A. 98:13183–13188.

Turner, B. C. 2001. Geographic distribution of neurospora spore killer strains and strains resistant to killing. Fungal Genet. Biol. 32:93–104.

Turner, B. C., and D. D. Perkins. 1979. Spore killer, a chromosomal factor in neurospora that kills meiotic products not containing it. Genetics 93:587–606.

Turner, E., D. J. Jacobson, and J. W. Taylor. 2011. Genetic Architecture of a Reinforced, Postmating, Reproductive Isolation Barrier between Neurospora Species Indicates Evolution via Natural Selection. PLoS Genet. 7:e1002204.

Villalta, C. F., D. J. Jacobson, and J. W. Taylor. 2009. Three new phylogenetic and biological Neurospora species: N. hispaniola, N. metzenbergii and N. perkinsii. Mycologia 101:777–789.

Vogan, A. A., S. L. Ament-Velásquez, A. Granger-Farbos, J. Svedberg, E. Bastiaans, A. J. Debets, V. Coustou, H. Yvanne, C. Clavé, S. J. Saupe, and H. Johannesson. 2019. Combinations of Spok genes create multiple meiotic drivers in Podospora. Elife 8:e46454.

Vogel, H. J. 1956. A Convenient Growth Medium for Neurospora crassa. Microbial Genetics Bulletin 13:42–47.

Welch, J. J. 2004. Accumulating Dobzhansky-Muller incompatibilities: reconciling theory and data. Evolution 58:1145–1156.

Westergaard, M., and H. K. Mitchell. 1947. Neurospora V. A Synthetic Medium Favoring Sexual Reproduction. Am. J. Bot. 34:573.

Wilkinson, G. S., and C. L. Fry. 2001. Meiotic drive alters sperm competitive ability in stalk-eyed flies. Proc. Biol. Sci. 268:2559–2564.

Wilkinson, G. S., J. G. Swallow, S. J. Christensen, and K. Madden. 2003. Phylogeography of sex ratio and multiple mating in stalk-eyed flies from southeast Asia. Genetica 117:37–46.

Zhang, L., T. Sun, F. Woldesellassie, H. Xiao, and Y. Tao. 2015. Sex ratio meiotic drive as a plausible evolutionary mechanism for hybrid male sterility. PLoS Genet. 11:e1005073.

