## Supplementary figures and images for "Meiotic drive is associated with sexual incompatibility in *Neurospora*"

### Supplementary Figure 1

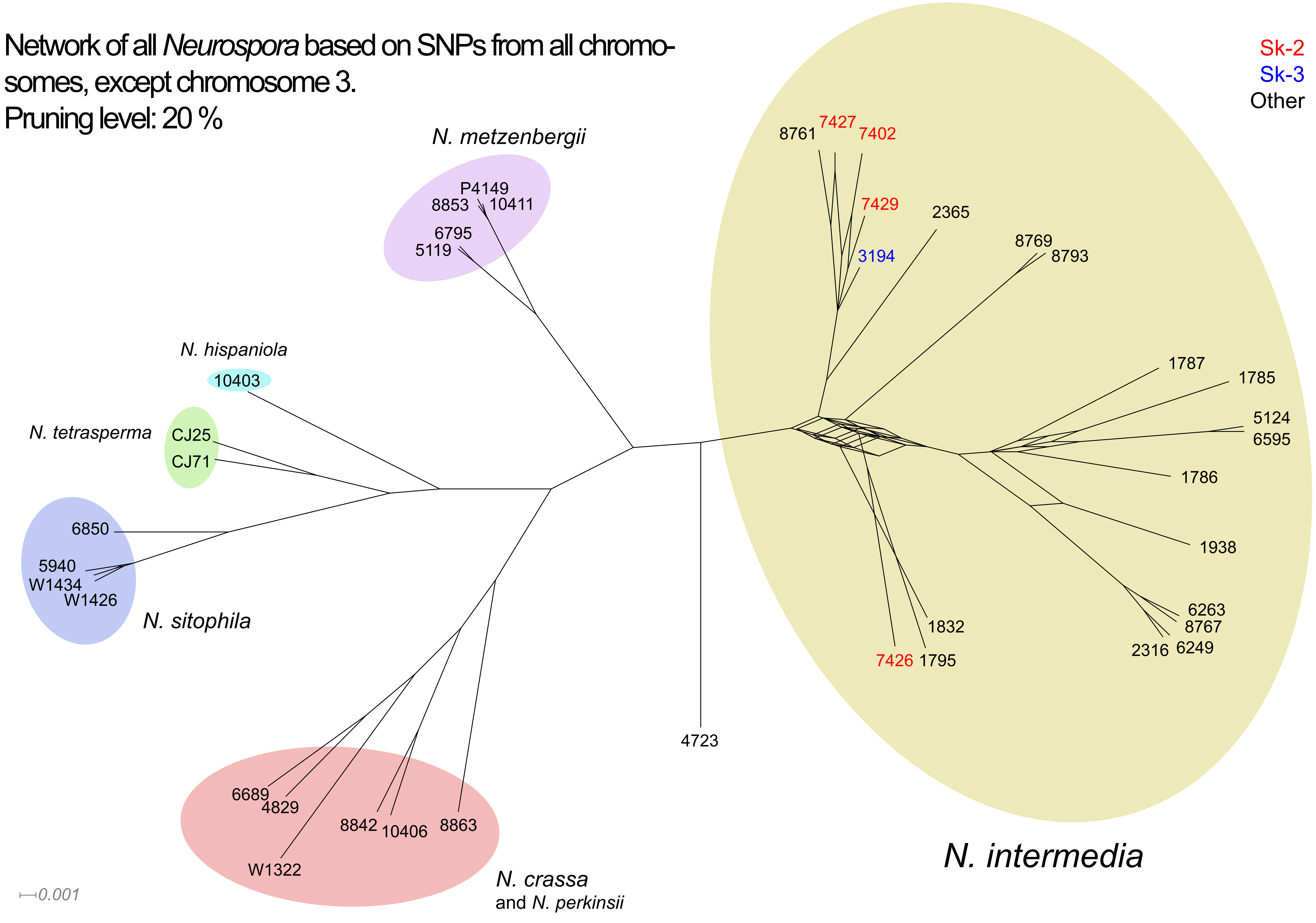
